# Universal pretreatment development for low-input proteomics using Lauryl Maltose Neopentyl Glycol

**DOI:** 10.1101/2023.08.10.552893

**Authors:** Ryo Konno, Masaki Ishikawa, Daisuke Nakajima, Yusuke Endo, Osamu Ohara, Yusuke Kawashima

**Affiliations:** Department of Applied Genomics, Kazusa DNA Research Institute, Kisarazu, Chiba 292-0818, Japan; Department of Frontier Research and Development, Kazusa DNA Research Institute, Kisarazu, Chiba 292-0818, Japan

**Keywords:** DMNG, LMNG, peptide loss, SCP, SP3

## Abstract

In recent years, the demand for low-input proteomics, most notably single-cell proteomics (SCP), has increased. In this study, we developed a Lauryl Maltose Neopentyl Glycol (LMNG)-assisted sample preparation (LASP) method that suppresses the loss of proteins and peptides in samples by adding LMNG, a surfactant, to the digested solution and removing the LMNG simply via reversed phase solid-phase extraction. The advantage of removing LMNG during sample preparation for general proteomic analysis is that it prevents mass spectrometry (MS) contamination. When the LASP method was applied to the low-input SP3 method and on-bead digestion in immunoprecipitation-MS, the recovery of the digested peptides was greatly improved. Furthermore, we established a simple and operationally easy sample preparation method for SCP based on the LASP method (scpLASP) and identified a median of 1,175 proteins from a single HEK239F cell using liquid chromatography (LC)-MS/MS with a throughput of 80 samples per day.

## Introduction

In samples with sufficient cell numbers for proteome analysis, the field has reached an era where a single-shot measurement can reveal more than 10,000 proteins (Kawashima *et al*, 2022; Muntel *et al*, 2019; Stewart *et al*, 2023). Proteome analysis is expected to make further contributions in the fields of biology and medicine. The next major challenge in proteome analysis is the active development of techniques for low-input samples such as single-cell proteomics (SCP) and spatial proteome analysis. However, the rates of protein and peptide loss increase as the amount of protein in the sample decreases, making single-cell and spatial proteomic analyses even more difficult. Techniques have been developed for sample preparation in nanoliter-scale liquid volumes, such as nanoPOTS (Williams *et al*, 2020; Woo *et al*, 2021; Zhu *et al*, 2018) and the OAD method (Li *et al*, 2018), which can minimize nonspecific adsorption for low-input samples; however, such methods are not general-purpose and require special equipment. Therefore, it is necessary to prevent protein and peptide loss using a simple method that is as versatile as possible, without the need for specialized equipment.

The major sample preparation steps for proteome analysis were 1) protein extraction, 2) reduction-alkylation, 3) enzymatic digestion, 4) desalting using a reversed phase solid-phase extraction (RP-SPE) column, 5) drying, and 6) redissolution. In addition, with the advent of the SP3 (Hughes *et al*, 2014; Hughes *et al*, 2019) and S-Trap methods (Thanou *et al*, 2023), versatile methods for removing surfactants and salts from lysates and enzymatic digestion are widely used. However, in low-input samples, proteins and peptides are lost during long incubations, such as during protein digestion, and upon transferring samples to a different tube. The use of surfactants is desirable to reduce these losses; however, surfactants are trapped in the RP-SPE column, elute with peptides, and cannot be easily removed. Known surfactants that do not interfere with peptide analysis by liquid chromatography-mass spectrometry (LC-MS/MS) include DDM (Liu *et al*, 2015; Nie *et al*, 2022; Tsai *et al*, 2021). However, for low-input samples, it is common to dry the eluted sample from an RP-SPE column, dissolve the peptides in a small amount of solvent, and analyze the concentrated peptides by LC-MS/MS. This method naturally increases the risk of MS contamination because the concentrated surfactant is analyzed with concentrated peptides. Depending on the surfactant type, methods to remove the surfactant after digestion by precipitation (Lin *et al*, 2013), phase transfer surfactants (Masuda *et al*, 2008), or decomposition of the surfactant (Chang *et al*, 2015; Mosen *et al*, 2021) have been developed; however, these treatments increase the number of work steps, resulting in various disadvantages, including increased labor, reduced reproducibility, and loss of peptides. Therefore, it is ideal for surfactants to be removed by a general process without any special requirements.

In this study, we searched for surfactants that can be removed using RP-SPE columns without any special treatment. Since it would be difficult to find a surfactant that does not adsorb onto an RP-SPE column because surfactants have both hydrophilic and hydrophobic groups, we attempted to find a surfactant with strong affinity to an RP-SPE column and to find conditions where only peptides elute without eluting the surfactant. The surfactants, including DDM, were obtained from sugar-type surfactants, which fall under a category known for longer elution times compared to peptides. A total of 12 surfactants were tested in the study. In addition, the use of low concentrations of surfactants for peptide redissolution was further investigated to reduce peptide loss due to the drying step after RP-SPE. Ultimately, we established a simple and easy pretreatment method for SCP and performed high-throughput single-cell proteomics (80 samples per day) using EVOSEP ONE.

## Results

### Development of a surfactant removal method using RP-SPE column for low-input proteomics

The surfactants listed in Table 1 were analyzed by LC-MS using a C18 column (**Fig 1A**). The results showed that Laury Maltose Neopentyl Glycol (LMNG) had a longer retention time than the other surfactants and a strong affinity for the RP column. The extraction conditions in RP-SPE (STAGE tip) were then examined for LMNG, DMNG, and DDM, which had longer retention times. In this study, a commercially available SDB-Stage tip, which is commonly used for desalting digested peptides, was used for RP-SPE. A 50–80% ACN solution is generally used as the eluent to ensure that the peptides are eluted. We trapped LMNG, DMNG, and DDM on the SDB-Stage tip and eluted sequentially with 30% ACN in 0.1% TFA, 40% ACN in 0.1% TFA, 50% ACN in 0.1% TFA, and 60% ACN in 0.1% TFA (**Fig 1B**). For DDM and DMNG, elution was achieved using 30% ACN in 0.1% TFA, necessitating a lower ACN concentration for their removal. Below 30% ACN, a risk of loss of peptides with a strong affinity for the RP carrier occurs. In contrast, LMNG was found not to be eluted at all in 30% ACN in 0.1% TFA and only slightly eluted in 40% ACN in 0.1% TFA. Furthermore, a detailed examination of the elution conditions for LMNG revealed that it did not elute at elution conditions below 38% ACN in 0.1% TFA (**Fig 1C**). Because the elution of SPE columns changed slightly depending on the room temperature and elution speed, 36% ACN-0.1% TFA was adopted as the condition under which LMNG was not eluted, with a generous margin. We then investigated whether 36% ACN in 0.1% TFA would be sufficient as an elution solvent for digested peptides (**Fig 1D**). As a result, the number of peptides identified did not change under the elution conditions of 50% ACN in 0.1% TFA and 36% ACN in 0.1% TFA, and a method was successfully established to remove LMNG without affecting peptide elution on a RP-SPE column. Because LMNG could be easily removed, we tested whether the addition of LMNG to the digestive solution would improve peptide recovery (**Fig 1E**). We treated 100 ng of HEK293 cell lysate (20 µL of 5 ng/µL protein concentration was used) with the generic SP3 method and confirmed that the addition of LMNG during digestion dramatically increased the number of peptides identified. Regarding the concentration of LMNG, 0.02% resulted in the highest number of identified peptides, making it the optimal concentration for the analysis. We call our LMNG-based method the LASP (<u>L</u>MNG-<u>A</u>ssisted <u>S</u>ample <u>P</u>reparation) method.

**Fig 1.**
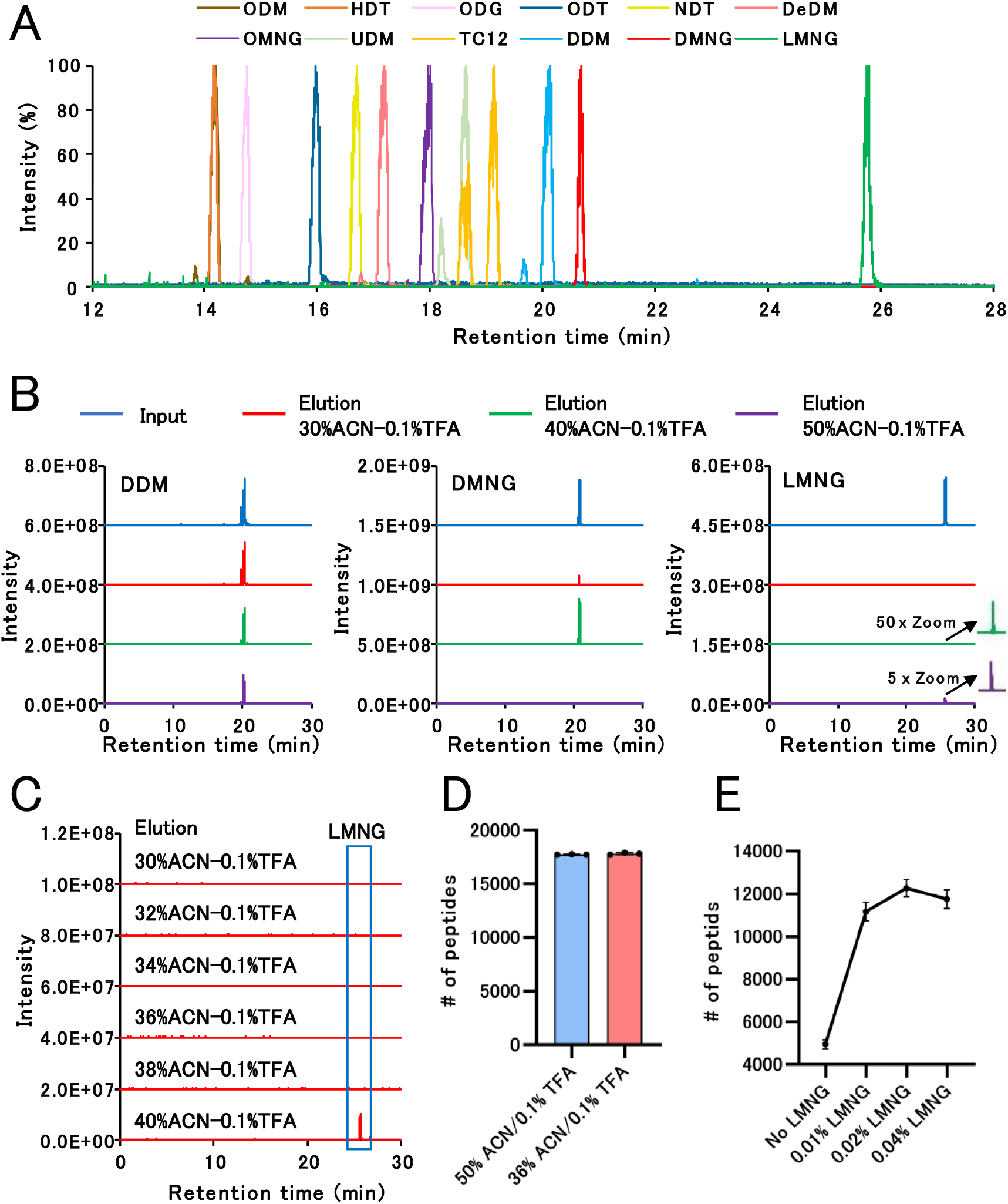
Development of LMNG removal method by RP-SPE. **(A)** MS chromatograms of the monoisotopic mass (single charge) of each surfactant. **(B)** Elution of LMNG, DMNG, and DDM from the SDB-Stage tip. LMNG, DMNG, and DDM were trapped in SDB-Stage tip and eluted sequentially with 30% ACN in 0.1% TFA, 40% ACN in 0.1% TFA, 50% ACN in 0.1% TFA, and 60% ACN in 0.1% TFA. Subsequently, eluted samples were analyzed by LC-MS. **(C)** Detailed elution conditions of LMNG from the SDB-Stage tips. LMNG was trapped in SDB-Stage tip and eluted with 30% ACN in 0.1% TFA, 32% ACN in 0.1% TFA, 34% ACN in 0.1% TFA, 36% ACN in 0.1% TFA, 38% ACN in 0.1% TFA, and 40% ACN in 0.1% TFA, in that order, and the eluted samples were analyzed by LC-MS. **(D)** Comparison of the identification number of peptides eluted from the SDB-Stage tip at 50% ACN in 0.1% TFA and 36% ACN in 0.1% TFA. Ten micrograms of HEK293 cell digest was dried from the tip after elution with each solvent. The dried peptide was then dissolved in 100 μL of 2% ACN in 0.1% TFA, and 1 μL was injected into DDA-LC–MS/MS. **(E)** Effects of LMNG on peptide recovery. HEK293 cell lysates (100 ng) were analyzed using the SP3 method. Subsequently, 50 mM Tris-HCl pH 8.0 (No LMNG) and 50 mM Tris-HCl pH 8.0 with 0.01% LMNG, 50 mM Tris-HCl pH 8.0 with 0.02% LMNG, and 50 mM Tris-HCl pH 8.0 with 0.04% LMNG were used as digestion solvents. LMNG was removed by SDB-Stage tip (elution: 36% ACN in 0.1% TFA), and peptides were dried. The dried peptide was then dissolved in 8 μL of 2% ACN in 0.1% TFA, and 4 μL was injected into DDA-LC– MS/MS.

**Table 1.** Surfactant list.

| Sugar-based surfactant | Abbreviation | Cas no | Molecular weight | CMC (mmol/l) | Company | CAT# |
| --- | --- | --- | --- | --- | --- | --- |
| n-Dodecyl-β-D-maltoside | DDM | 69227-93-6 | 510.62 | 0.17 | FUJIFILM Wako Pure Chemical Corp. | 347-06163 |
| n-Undecyl-β-D-maltoside | UDM | 253678-67-0 | 496.59 | 0.55 | Merck | 94206 |
| n-Decyl-β-D-maltoside | DeDM | 82494-09-5 | 482.57 | 1.8 | FUJIFILM Wako Pure Chemical Corp. | 349-08041 |
| n-Octyl-β-D-maltoside | ODM | 82494-08-4 | 454.51 | 23.4 | FUJIFILM Wako Pure Chemical Corp. | 340-90283 |
| n-Nonyl-β-D-thiomaltoside | NDT | 148565-55-3 | 484.6 | 2.4 | FUJIFILM Wako Pure Chemical Corp. | 343-06861 |
| n-Octyl-β-D-glucoside | ODG | 29836-26-8 | 292.37 | 25 | FUJIFILM Wako Pure Chemical Corp. | 346-05033 |
| n-Heptyl-β-D-thioglucoside | HDT | 85618-20-8 | 294.41 | 30 | FUJIFILM Wako Pure Chemical Corp. | 346-05371 |
| n-Octyl-β-D-thioglucoside | ODT | 85618-21-9 | 308.44 | 9 | FUJIFILM Wako Pure Chemical Corp. | 349-05361 |
| Decyl Maltose Neopentyl Glycol | DMNG | 1257852-99-5 | 949.08 | 0.036 | Anatrace | NG322 |
| Lauryl Maltose Neopentyl Glycol | LMNG | 1257852-96-2 | 1005.19 | 0.01 | Anatrace | NG310 |
| Octyl Glucose Neopentyl Glycol | OMNG | 1257853-32-9 | 568.69 | 1.02 | Anatrace | NG311 |
| Trehalose C12 | TC12 | 64622-91-9 | 524.6 | 0.15 | FUJIFILM Wako Pure Chemical Corp. | 340-09051 |

### Resolubilization of dried peptides

To clarify the changes in the recovery of dried peptides by surfactants, we prepared small amounts (500 ng) of dried tryptic peptides, which are prone to peptide loss. The dried peptides were dissolved in distilled water, 2% ACN in 0.1% TFA, and 12 surfactants (concentration: 0.04%) for DDA-LC-MS/MS analysis (**Fig 2A**). The number of identified peptides increased with the use of surfactants. In addition, to identify a surfactant that minimizes MS contamination and dissolves peptides, we tested the effect of dissolving peptides at lower concentrations using nine surfactants, for which we were able to identify more than 18,000 peptides (**Fig 2B**). The number of peptides identified in NDT, ODT, and DeDM remarkably decreased at lower concentrations. At 0.01% NDT, ODT, and DeDM, the number of identified peptides was considerably reduced. Other surfactants showed a marked decreasing trend starting at 0.005%. Therefore, validation was performed on the top five surfactants (DDM, UDM, DMNG, OMNG, and TC12) with the highest number of peptides identified at 0.01% (**Fig 2C**). Although there were no significant changes due to the surfactants selected, all of these surfactants were shown to be effective in peptide recovery, even at concentrations as low as 0.01%. Additionally, we investigated the length of peptides identified by each solution and found that the difference in the number of peptides identified between 2% ACN in 0.1% TFA, and these surfactants, increased as the peptides became longer (**Fig 2D**). Thus, the use of these surfactants improved the recovery of longer peptides. All five surfactants were effective in dissolving low concentrations of dried peptides; however, we decided to use DMNG, which has the latest retention time that differs the most from the retention time of the peptides.

**Fig 2.**
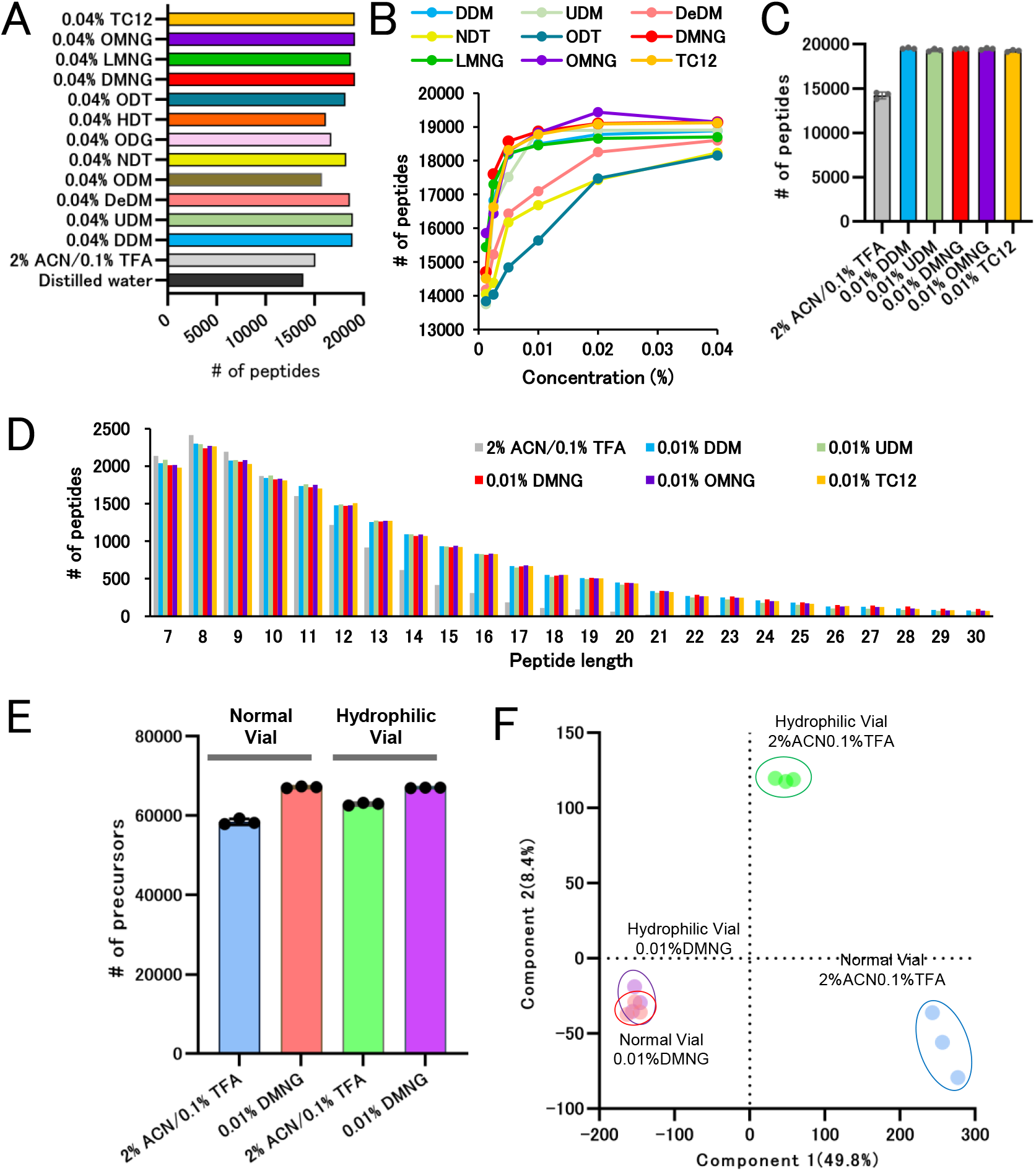
The recovery of dried tryptic peptides by surfactants. **(A)** Number of peptides identified from dried peptides dissolved in 12 surfactants at 0.04%, 2% ACN in 0.1% TFA, and distilled water. The dried peptide (500 ng) in a 1.5 mL tube was dissolved in 25 μL of each solvent, transferred to a normal PP vial, and 1 μL was injected into DDA-LC–MS/MS (the same treatment was carried out for (**B**) and (**C**)). **(B)** Number of peptides identified from dried peptides dissolved in nine selected surfactants at 0.00125–0.04%. **(C)** Number of peptides identified from dried peptides dissolved in five selected surfactants at 0.01% and 2% ACN in 0.1% TFA (n=3). **(D)** Distribution of amino acid lengths of peptides identified from dried peptides dissolved in each solvent seen in (**C**). **(E)** Number of precursors identified from dried peptides in normal PP vials or hydrophilic vials dissolved in 2% ACN–0.1% TFA or 0.01% DMNG. The dried peptide (500 ng) in each vial was dissolved in 25 μL of each solvent, and 1 μL was injected into the DIA-LC-MS/MS. **(F)** Principal component analysis of precursor intensity of the results in (**E**).

Next, we measured and compared the recovery of 500 ng of dried tryptic peptide in hydrophilic-coated vials and normal PP vials. This was dissolved in 2% ACN in 0.1% TFA or 0.01% DMNG (four-combinations in total) for DIA–LC–MS/MS quantitative analysis. We identified more precursors in the hydrophilic-coated vials dissolved in 2% ACN in 0.1% TFA and no change in the hydrophilic-coated vials dissolved in 0.01% DMNG (**Fig 2E**). More precursors were identified in the dissolution with 0.01% DMNG than in the dissolution with 2% ACN in 0.1% TFA, regardless of the vial. PCA analysis based on precursor intensity also showed no differences between vials for DMNG, but there were differences between vials for 2% ACN in 0.1% TFA, with the 2% ACN in 0.1% TFA-hydrophilic vial group being closer to the group dissolved in DMNG (**Fig 2F**). These results demonstrated improved dried peptide recovery with the use of hydrophilic vials for the dissolution with 2% ACN in 0.1% TFA. In contrast, upon dissolving in 0.01% DMNG, no differences between vials were observed. Other than the difference between vials, the use of 0.01% DMNG substantially contributed to the amount of recovered peptides. Therefore, the normal vials were sufficient for use with solutions containing 0.01% DMNG.

### Combined effect of LASP and DMNG dissolution methods

To minimize peptide loss, we tried both adding 0.02% LMNG to the digestion solvent and using 0.01% DMNG to dissolve the dried peptides. In **Fig 3A** and **B**, 200 µL of 0.5 ng/µL HEK293 cell lysate (protein amount: 100 ng) was used as starting sample and treated using the SP3 method. The SP3 method faces challenges when processing samples with such low concentrations. The addition of LMNG to the digestion solvent markedly increased the number of identified precursors and proteins, and dissolving dried peptides in DMNG further increased these numbers (**Fig 3A**). In addition, in samples without LMNG, the quantitative values of the individual precursors varied remarkably, resulting in low Pearson’s r values (**Fig 3B**). In contrast, the LMNG samples exhibited higher Pearson’s r values. In the case of low-concentration samples, such as those used in this study, the addition of LMNG not only improved the number of identified proteins and precursors, but also affected the reproducibility of sample preparation. Next, we investigated the effect of adding LMNG during on-bead digestion using co-immunoprecipitation-mass spectrometry (coIP-MS) as well as the effect of dissolving dried peptides in DMNG (**Fig 3C–E**). Here, coIP-MS was performed using an anti-RELA antibody. As in the SP3 experiment, the addition of LMNG markedly improved the identification numbers of the precursors and proteins, and dissolving the dried peptides in DMNG further increased these numbers (**Fig 3C**). Although the coIP-MS study maintained the reproducibility of precursor quantitation values for individual groups, the LMNG-treated samples had higher Pearson’s r values regardless of DMNG, and LMNG contributed extensively to the efficient and stable recovery of peptides in this experiment (**Fig 3D**). The recovery of RELA, the antibody target, and its known interactors, NFKB1, IKBKB, NFKBIA, NFKBIB, and NFKBIE, was greatly improved by LMNG and further improved upon lysis with DMNG (**Fig 3E**). These results confirmed that both the LASP and peptide dissolution methods with DMNG have beneficial effects on coIP-MS. Improvements in the number of identified phosphopeptides and biotinylated peptides were achieved by adding LMNG to the elution solvent in the phosphopeptide and biotinylated peptide enrichment experiments using immobilized metal ion affinity chromatography and streptavidin beads, respectively (**Fig 4A and B**). Because LMNG can be easily removed by RP-SPE columns, the LASP method is expected to have a wide range of applications, not only for reducing peptide loss during digestion, but also for affinity purification of peptides.

**Fig 3.**
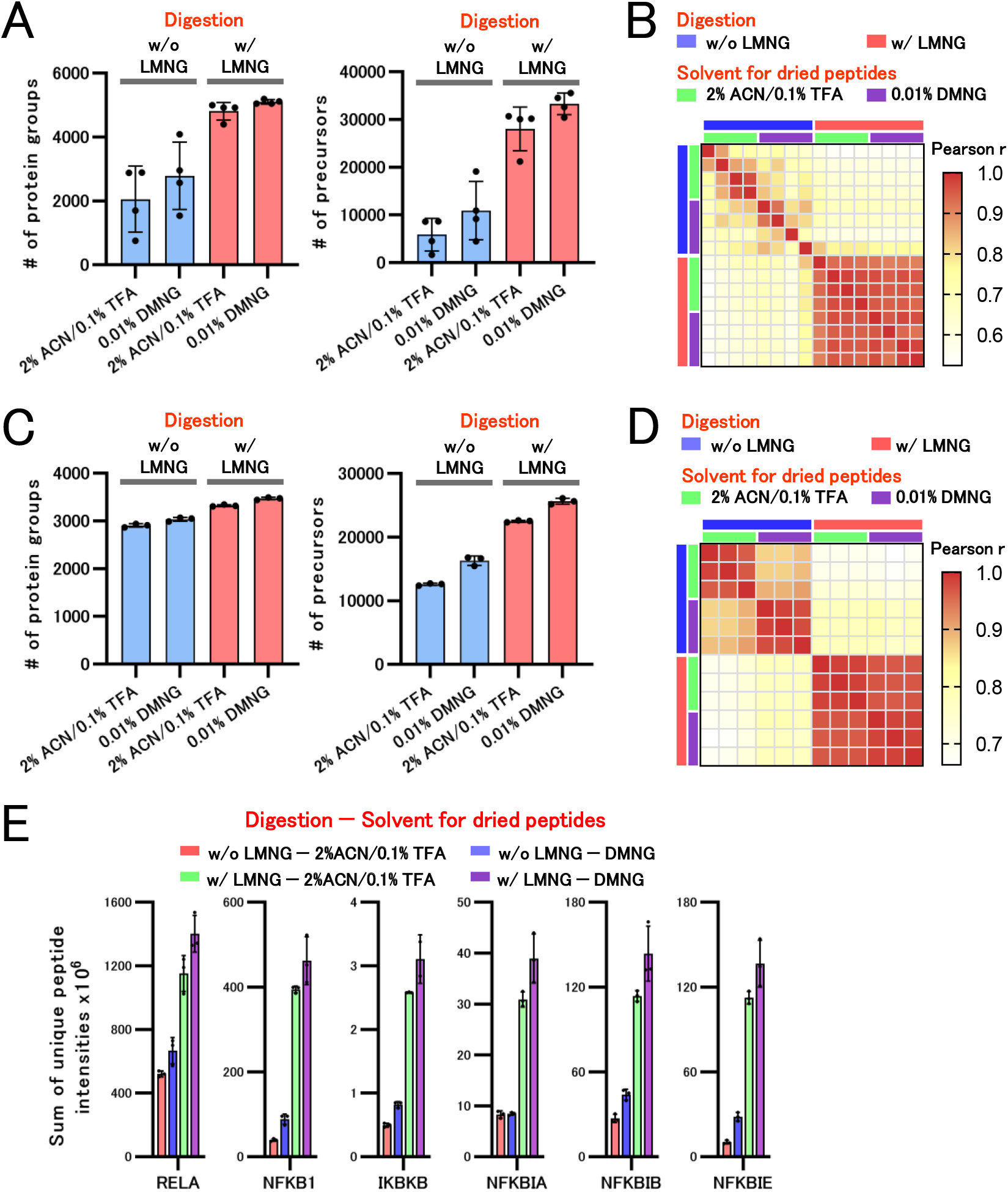
Combined effect of LASP and DMNG dissolution methods for SP3 and coIP methods. **(A)** The number of proteins and precursors identified with and without the addition of LMNG during digestion by the SP3 method and by dissolving dried peptides. 200 µL of 0.5 ng/µL HEK293 cell lysate was treated using the SP3 method. 50 mM Tris-HCl pH 8.0 with or without 0.02% LMNG was used as the digestion solvent. LMNG was removed using an SDB-Stage tip, and the peptides were dried. The dried peptide was then dissolved in 8 μL of 2% ACN in 0.1% TFA or 0.01% DMNG, and 4 μL was injected into DIA-LC–MS/MS. **(B)** Pearson correlation coefficient heatmap with hierarchical clustering of precursor intensity in HEK293 cell proteome analysis. **(C)** The number of proteins and precursors identified with and without the addition of LMNG during on-bead digestion by the coIP-MS method and by dissolving dried peptides. The coIP was performed using an anti-RALA antibody. 50 mM Tris-HCl pH 8.0 with or without 0.02% LMNG was used as the digestion solvent. LMNG was removed using an SDB-Stage tip, and the peptides were dried. The dried peptide was then dissolved in 10 μL of 2% ACN in 0.1% TFA or 0.01% DMNG, and 1 μL was injected into DIA-LC–MS/MS. **(D)** Pearson correlation coefficient heatmap with hierarchical clustering of precursor intensities of coIP-MS. **(E)** The sum of the unique peptide intensities of RELA and renowned RELA interactors in the identified proteins.

**Fig 4.**
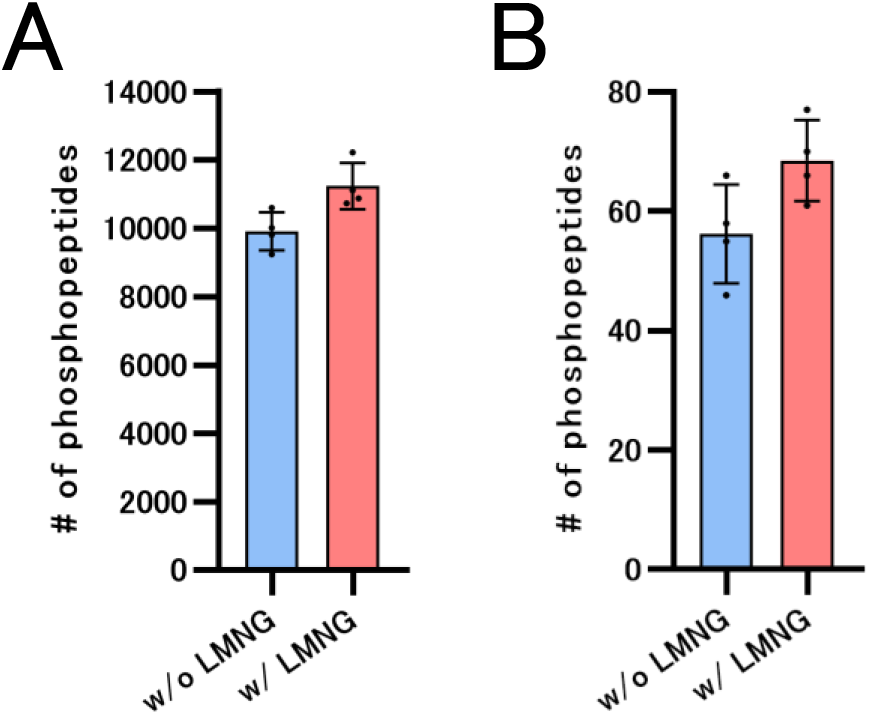
Effect of LMNG-added elution solvent on affinity purification of peptides. **(A)** Comparison of phosphopeptides identified with and without LMNG addition during the elution of Fe-NTA magnetic IMAC beads. Phosphopeptides were enriched from 100 µg of HEK293 cell digests. **(B)** Comparison of biotinylated BSA peptides identified with and without LMNG addition during streptavidin bead elution Biotinylated BSA peptides were enriched from a mixture of 10 ng of biotinylated BSA digest and 5 µg of K562 cell digest.

### Benefits of the LASP method in high-throughput proteomics using EVOSEP ONE

The EVOSEP ONE is a well-known LC instrument for high-throughput proteomics and is used worldwide for the proteomic analysis of hundreds to thousands of samples, such as clinical samples. The EVOSEP ONE allows for LC analysis by trapping the sample in an RP-SPE column [Evotip pure (C18)] and setting it in the EVOSEP ONE. For EVOSEP ONE, the manufacturer has already prepared an optimized method for peptide analysis that does not allow for fine gradient changes. First, we examined whether LMNG was eluted from the EVOSEP ONE. In this study, a method with a throughput of 80 samples per day (SPD) was utilized for all analyses using the EVOSEP ONE. LMNG and, as controls, DDM and DMNG were also trapped in Evotip Pure and measured by MS after separation using an EVOSEP ONE (**Fig 5A**). Both DDM and DMNG were detected at the end of the retention time, indicating that they were eluted from Evotip Pure, but not from LMNG. When EVOSEP ONE was used, 100 ng of HEK293 cell lysate at the same low concentration as shown in **Fig 3** was digested with and without LMNG and analyzed by LC-MS/MS (**Fig 5B and C**). More precursors and proteins were identified in LMNG. In addition, even when extracellular vesicles were purified from serum samples and analyzed by LC-MS/MS using EVOSEP ONE, the addition of LMNG during digestion improved the number of identified precursors and proteins and the recovery of typical exosome markers, such as CD9, CD63, and CD81 (**Fig 5D and E**). The reproducibility of the quantitative precursor values was further improved upon the addition of LMNG in both experiments (**Fig 5C and F**). LMNG was also found to be highly compatible with high-throughput proteomic analysis using EVOSEP ONE.

**Fig 5.**
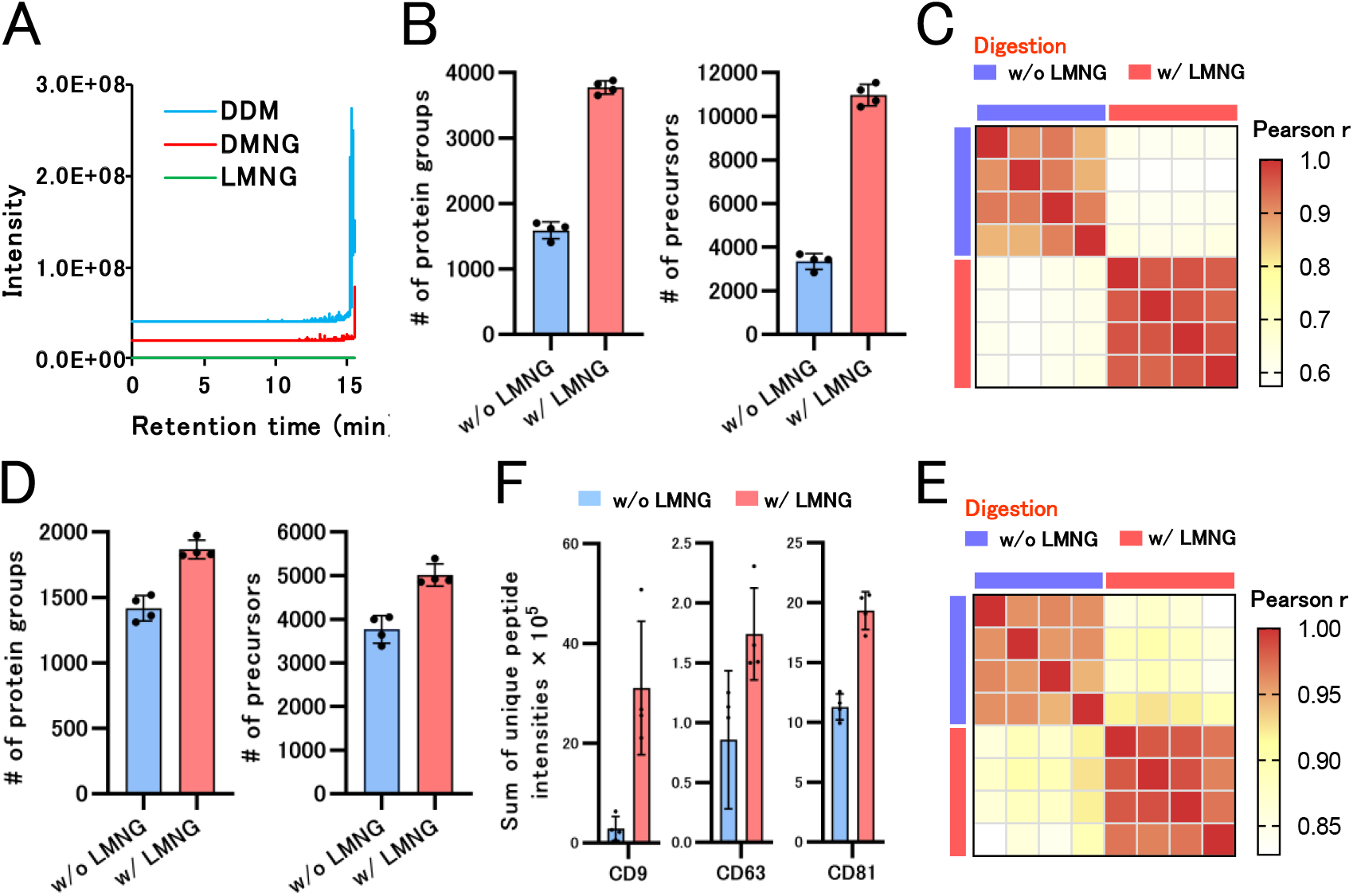
LASP method using Evosep One system. **(A)** Elution of LMNG, DMNG, and DDM using EvoSep-One system. LMNG, DMNG, and DDM were trapped in an Evosep Pure tip and set on the Evosep ONE. The Whisper 80 SPD method was used for the LC gradient of Evosep ONE (the same method was used in (**B**) and (**D**)). **(B)** The number of proteins and precursors identified with and without LMNG addition during the digestion of HEK293 cell lysates. 200 µL of 0.5 ng/µL HEK293 cell lysate was treated with the SP3 method. 50 mM Tris-HCl pH 8.0 with or without 0.02% LMNG was used as the digestion solvent. **(C)** Pearson correlation coefficient heatmap with hierarchical clustering of precursor intensities in HEK293 cell proteome analysis. **(D)** Number of proteins and precursors identified with and without LMNG addition during serum EVs digestion. EVs enriched with 100 µL of serum were treated using the SP3 method. **(E)** Sum of the unique peptide intensities of CD9, CD63, and CD81, which are known exosome markers. **(F)** Pearson correlation coefficient heatmap with hierarchical clustering of precursor intensities in serum EV proteome analysis.

### The Challenge of Label-Free SCP

SCP requires high throughput to analyze an extremely large number of cells. Therefore, we attempted to establish a system using EVOSEP ONE with the 80 SPD method for SCP. For the analytical conditions of MS, we tested the parameters of DIA-MS, thereby allowing for the observation of a large number of proteins, even in trace samples, in the gradient of the 80 SPD method. MS/MS analysis of the m/z 550-670, m/z 600-720, and m/z 650-770 regions with an MS2 resolution of 60 K and an isolation window width of 15 Th in the first screening revealed a higher number of protein identifications (**Fig 6A**). Furthermore, when the three methods were validated, the method targeting m/z600-720 identified more proteins, albeit only slightly (**Fig 6B**). All methods showed high protein quantification reproducibility (**Fig 6C**). Based on these results, we adopted MS/MS at 15 Th intervals for m/z600-720 at an MS2 resolution of 60 K as the DIA method for the SCP.

**Fig 6.**
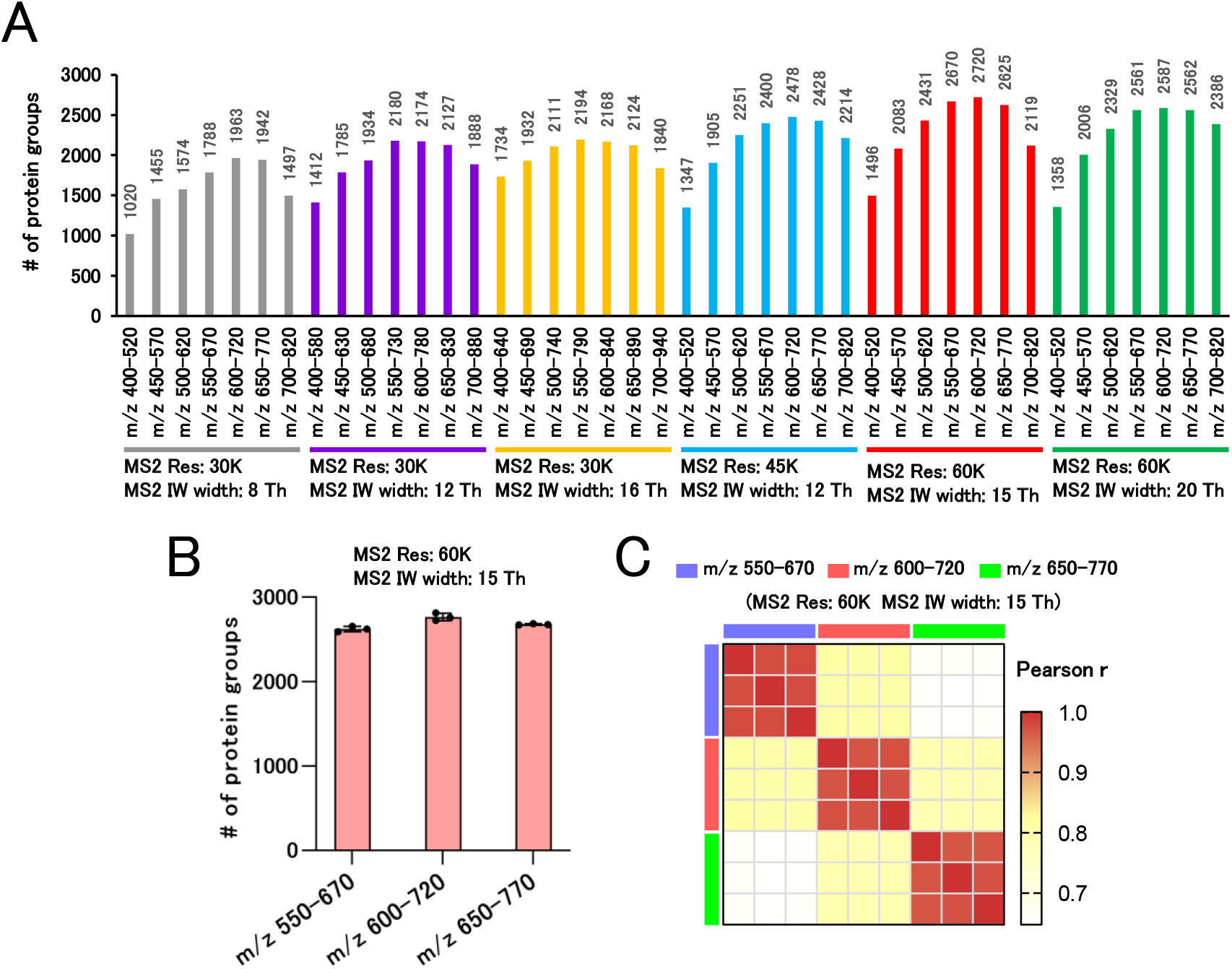
Investigation of DIA-MS parameters for high-throughput few-cell proteomics analysis. **(A)** Comparison of protein groups identified using various DIA-MS parameters. One nanogram of the HEK293 cell digest was trapped in an Evosep Pure tip and placed on Evosep ONE. The Whisper 80 SPD method was used for the LC gradient of Evosep ONE (the same method was used in (**B**)). **(B)** Validation of the method for MS/MS analysis of the m/z 550-670, m/z 600-720, and m/z 650-770 regions with an MS2 resolution of 60 K and an isolation window width of 15 Th. One nanogram of HEK293 cell digest was measured in triplicate for each method. **(C)** Pearson’s correlation coefficient heatmap with hierarchical clustering of precursor intensity in the proteome analysis results of 1 ng of HEK293 cell digest.

For sample preparation, we established the LASP method for scp (scpLASP), which is 96-well based and can process multiple samples simultaneously (**Fig 7A**). The scpLASP method uses a starting volume of 10 µL, making it easy to work with both manually and automatically. The scpLASP method was also evaluated with and without the addition of LMNG to the digestion solvent (**Fig 7B**). Both 1 cell and 10 cells analyses of HEK293 cells showed that the addition of LMNG remarkably increased the number of proteins identified. In scpLASP, protein identification analysis using the *in silico* spectral library generated by DIA-NN identified a median of 606 and 2,650 proteins in 1 and 10 cells, respectively. Furthermore, the results of the 10-cells analysis processed by the scpLASP method were converted to a spectral library. Using that library to analyze the MS data of one cell, a median of 1,175 proteins was successfully identified. This was effective in creating a spectral library with bulk samples in advance to perform label-free SCP.

**Fig 7.**
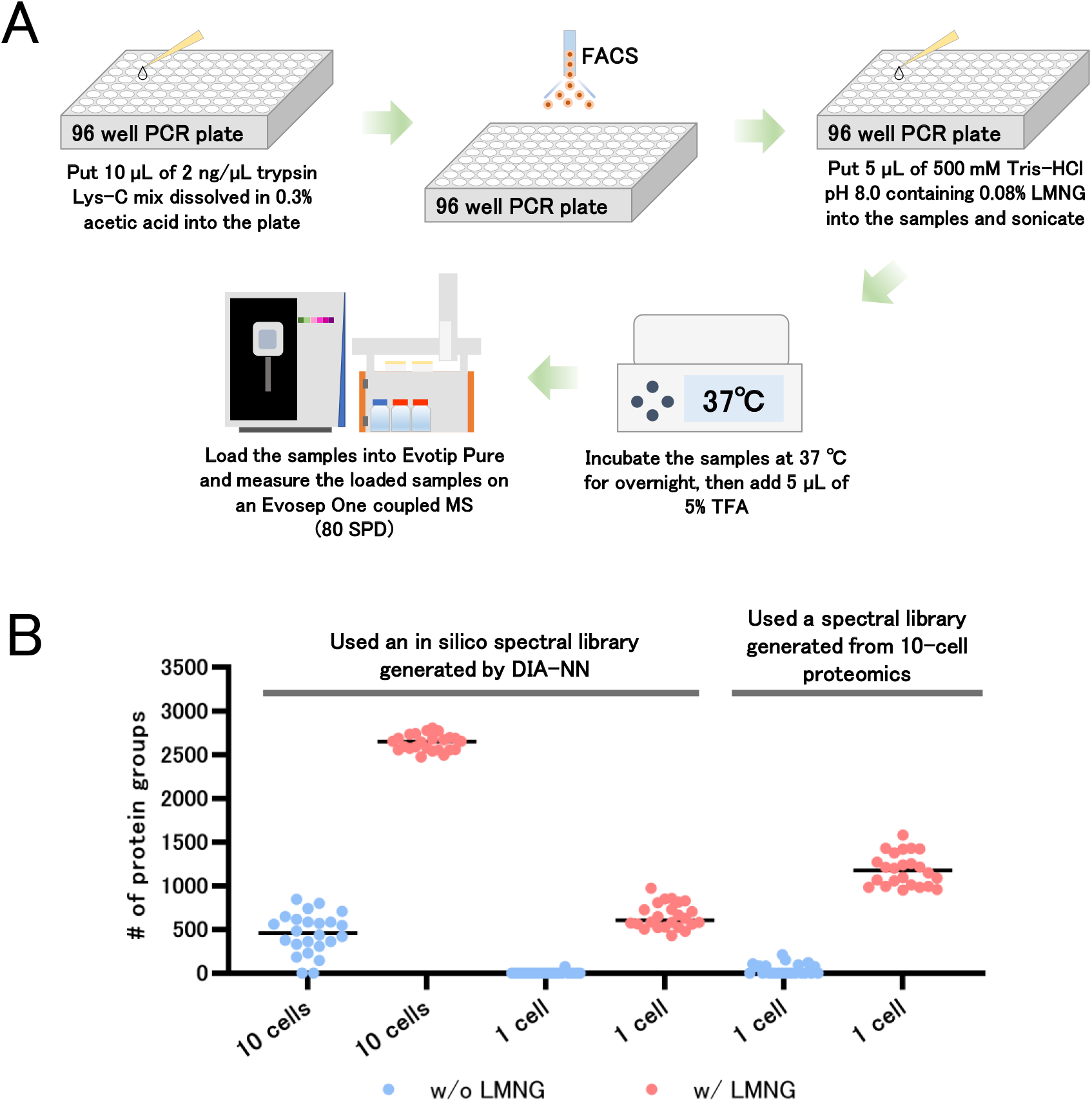
Label-free SCP by scpLASP method. **(A)** Workflow of the scpLASP method. We added 10 µL of 2 ng/µL trypsin Lys-C mix in 0.3% acetic acid to a 96-well plate and sorted cells with FASC. Then, we added 5 µL of 500 mM Tris-HCl pH 8.0 containing 0.08% LMNG, followed by sonication and overnight incubation at 37°C. After that, we added 5 µL of 5% TFA to the digested sample and loaded it into Evotip for analysis on an Evosep One coupled MS with a throughput of 80 SPD. **(B)** Single and ten cells proteomics with and without LMNG addition during digestion. The DIA-MS data of single and ten cells were first analyzed by the DIA-NN using an *in silico* spectral library. Single-cell DIA-MS data were then analyzed by DIA-NN using a spectral library generated from protein identification information obtained from the proteomics of 10 cells.

## Discussion

We developed the LASP method to suppress peptide loss by LMNG for easy low-input proteome analysis. Regarding the use of surfactants in low-input proteome analysis, Shi *et al*. (2021) reported reduced peptide loss with the addition of DDM (Martin *et al*, 2021; Tsai *et al*, 2021). DDM is highly effective because it does not interfere with peptide analysis during LC-MS/MS. However, DDM cannot be easily removed, and when analyzing by LC-MS, depending on the concentration of DDM at the start and the volume of liquid, a large amount of DDM is injected into the LC-MS, increasing the risk of MS contamination (in the case of scp, 2µL of 0.1% DDM was used for cell lysis). However, if the volume of liquid at the start is markedly reduced, handling becomes difficult, and the method cannot be used universally. From this perspective, LMNG has a considerable advantage in that it can be easily removed with RP-SPE, even if the amount used increases, making it a simple and versatile method. We also used a surfactant to re-dissolve dried peptides. However, we used a low concentration of 0.01%, and the volume of liquid dissolved was controlled to reduce contamination of the LC-MS by the surfactant. Even if the peptides are dissolved within a few microliters, which makes handling more difficult, the impact on the overall protocol is small because it is the final step in the sample preparation process.

To investigate the dissolution of dried peptides, we identified surfactants (DMNG, DDM, UDM, OMNG, and TC12) with high peptide recovery at low concentrations, which can be used for peptide analysis in C18-LC-MS/MS. For DDM, Nie et al. reported improved peptide recovery similar to our results (Nie *et al*, 2022). The effects of DMNG, UDM, OMNG, and TC12 were, for the first time, investigated. Although LMNG was also effective in recovering dried peptides, the affinity of LMNG for C18 was so strong that the risk of carryover was high and LMNG was not suitable for continuous analysis. However, LMNG may be suitable for protein lysis in top-down proteomic analyses. Because the C4 and C8 columns were used in top-down proteome analysis, the risk of LMNG carryover was lower than that for C18. Moreover, in RP-HPLC, proteins tend to elute at later retention times than peptides, which raises the concern of potential overlap between protein elution times and surfactants like DMNG, DDM, UDM, OMNG, and TC12, commonly used in peptide analysis. Therefore, the use of surfactants with extremely late elution times, such as LMNG, offers an advantage in this context. The late retention time characteristic of LMNG has both advantages and disadvantages, but we believe that it can be widely applied if it is exploited. After 2 months of measuring various digests dissolved in DMNG for MS contamination, there was no significant change in the number of proteins identified in the HEK293 digests, which were regularly measured for MS quality control. The use of low concentrations of DMNG (0.01%) did not present significant problems with LC-MS. Hydrophilic coating of the LC vials was not necessary as long as the peptides were dissolved in DMNG and the normal PP vials were sufficient. Hydrophilic-coated vials and tubes are expensive, whereas dissolution with surfactants such as DMNG is inexpensive because high numbers of peptides can be recovered even with normal PP vials and tubes.

In SCP, hundreds to thousands of analyses are necessary, requiring high-throughput analysis of samples that are orders of magnitude larger than those used in the past. Therefore, the risk of MS contamination should be eliminated as much as possible. The combined LASP and EVOSEP system is an excellent method for peptide loss, sensitivity, throughput, and MS contamination. Although SCP sample preparation is generally performed with a small volume of liquid to reduce protein and peptide loss, we established the scpLASP method, which eliminates delicate handling of the sample as much as possible. A typical SCP requires careful setting up of the wells such that the sorted cells from the FACS are sent to the bottom of the wells, and the cells are placed in a few microliters of solvent placed at the bottom of the wells (Brunner *et al*, 2022; Petrosius *et al*, 2022; Tsai *et al*, 2021). The scpLASP method encompassed 10 µL in the bottom of the well to allow for easy collection of sorted cells. The presence of 10 µL of liquid from the start lowers the risk of the sample drying out, facilitates the addition of reagents and agitation, and simplifies the overall handling of the sample preparation. A higher liquid volume does not represent a major problem in our method because the surfactant added to reduce protein and peptide loss can be removed at the end of the Evotip Pure. In addition, 10 µL of 2 ng/µL trypsin Lys-C in 0.3% acetic acid is dispensed in 96 PCR well plates and stocked at -80°C for less preparation during FACS. Enzymatic digestion was initiated by adding 500 mM Tris-HCl pH 8.0 with 0.08% LMNG to the sorted cell samples, and peptide loss was also reduced. Thus, our scpLASP method is simplified, and we believe that it is an effective method not only for SCP, but also for spatial proteomics that processes multiple samples. Thus, we believe that the scpLASP method will become the standard method for ultra-low-input sample preparation, such as in SCP and spatial proteomics, owing to its simplicity and low risk of MS contamination.

Our SCP results showed that the combination of a simple pretreatment and 80 SPD LC-MS/MS allowed for the identification and quantification of a median of 1,175 proteins from a single HEK293 cell without enhancing the observed protein count by MBR, indicating that our total analysis system was also superior. However, there are more optimal conditions for LC column and MS equipment. For LC column, micropillar array columns suitable for low-input samples have been developed and are commercially available, albeit expensive. Compared to the 50 cm packed bed column, the 50 cm pillar column identified more than 1.5 times of proteins in the LC-MS/MS analysis of 10 ng of the HeLa digest (Stadlmann *et al*, 2019). In label-free SCP, although the LC-MS/MS throughput was 40 SPD, an average of 1,543 and 1,404 proteins were quantified in a single HeLa cell and a single K562 cell, respectively, using a 5.5 cm micro-pillar array column (Matzinger *et al*, 2023), and approximately 2,000 proteins were quantified from a single HEK293 cell using a 50 cm micropillar array column (Petrosius *et al*, 2022). MS data acquisition and data analysis methods were developed for SCP in these studies; however, the benefits of using micropillar array columns were considered significant. Furthermore, in June 2023, Orbitrap Astral (Thermo Fisher Scientific) was released as an ultrasensitive MS, and surprisingly, upon using the Orbitrap Astral with a 50 cm micropillar array column, approximately 2,500 proteins were identified from a single HEK293 cell at an LC-MS/MS throughput of 80 SPD (Petrosius *et al*, 2023). Bruker Daltonics Inc. simultaneously released timsTOF Ultra, an ultrasensitive MS, which holds tremendous boon to SCP. We believe that by combining these new techniques with the scpLASP method, a high-level SCP can be easily achieved.

## Materials and Methods

### LC-MS measurement of surfactants

The surfactants (Table 1) were directly injected onto a 75 μm × 12 cm nanoLC column (Nikkyo Technos Co., Ltd., Tokyo, Japan) at 50°C and then separated with a 30-min gradient (A = 0.1% FA in water, B = 0.1% FA in 80% ACN) consisting of 0 min 5% B, 25 min 95% B, 30 min 95% B at a flow rate of 300 nL/min using an UltiMate 3000 RSLCnano LC system (Thermo Fisher Scientific, Waltham, MA, USA). The elution from the column was analyzed using a Q Exactive HF-X (Thermo Fisher Scientific) with DDA. MS1 spectra were collected in the range of 100– 1,500 m/z at a 60,000 resolution to set an auto-gain control (AGC) targets of 1 × 10^6^ and a maximum injection time of 119 ms. The MS chromatograms of the monoisotopic mass (single charge) of each surfactant were obtained.

### Elution conditions for SDB-STAGE tip of LMNG, DMNG, and DDM

The SDB-STAGE tip was washed with 25 μL of 80% ACN in 0.1% TFA, followed by equilibration using 50 μL of 3% ACN in 0.1% TFA. Then, 10 μL of 0.001% LMNG, DMNG, or DDM was loaded onto tip, washed using 80 μL of 3% ACN in 0.1% TFA, and eluted stepwise with 50 μL of 30% ACN in 0.1% TFA, 40% ACN in 0.1% TFA, and 50% ACN in 0.1% TFA. An additional LMNG sample was eluted with 50 μL of 30% ACN in 0.1% TFA, 32% ACN in 0.1% TFA, 34% ACN in 0.1% TFA, 36% ACN in 0.1% TFA, 38% ACN in 0.1% TFA, and 40% ACN in 0.1% TFA, respectively. The eluate was dried using a centrifugal evaporator (miVac Duo Concentrator; Genevac Ltd., Ipswich, UK). The dried sample was redissolved in 200 μL of H_2_O and transferred to a normal vial (CAT# C5000-97, Thermo Fisher Scientific).

### Dried peptide mixtures

100 μg of K562 cell tryptic digest (CAT# V6951, Promega, Madison, WI, USA) was adjusted to 50 ng/μL with 50% ACN containing 0.1% TFA and dispensed in 10 μL each into normal tubes (1.5-mL safe-lock tube, CAT# 0030120086, Eppendorf, Hamburg; Germany), normal polypropylene (PP) vials (CAT# C5000-97, Thermo Fisher Scientific), or hydrophilic vials (ProteoSave vial, CAT# 11-19-1021-10, AMR Inc., Tokyo, Japan). These samples were dried in a centrifugal evaporator (miVac Duo concentrator, Genevac Ltd., Ipswich, UK) and redissolved in 25 μL of distilled water, 2% ACN containing 0.1% TFA, or 0.00125-0.04% surfactants in distilled water (Table 1).

### Protein extraction from HEK293T cells

Proteins from the HEK293T cells were extracted in 100 mM Tris-HCl (pH 8.0) and 20 mM NaCl containing 4% sodium dodecyl sulfate (SDS) by sonication in Bioruptor II (CosmoBio, Tokyo, Japan) for 10 min. Protein concentration in the protein extract was determined using a BCA protein assay kit (CAT# 23225, Thermo Fisher Scientific) and adjusted to 500 ng, 5 or 0.5 ng/μL with 100 mM Tris-HCl (pH 8.0), and 20 mM NaCl containing 4% SDS.

### Purification of extracellular vesicle (EVs) from human serum

Human serum EVs were purified using the Tim4-phosphatidylserine (PS) affinity method combined with the MagCapture Exosome Isolation Kit PS Ver.2 (Wako Pure Chemical, Osaka, Japan) and Maelstrom 8 Autostage (Taiwan Advanced Nanotech Inc., Taoyuan, Taiwan). Briefly, 150 μL of pooled healthy human serum was centrifuged at 3,000 × *g* and 4°C for 20 min, and then, 100 μL of supernatants were collected in different tubes. Subsequently, 300 μL of TBS and 1 μL of Exosome Binding Enhancer (provided with the kit) were added to the supernatants and gently mixed. To prepare the beads for Tim4-PS affinity method, 30 μL of Exosome Capture Beads (provided with the kit) was washed once with 250 μL of Exosome Immobilizing/Washing Buffer (provided with the kit). Then, the beads were mixed in 10 μL of Biotin-labeled Exosome Capture (diluted by 250 μL of Exosome Immobilizing/Washing Buffer) and agitated at 1,000 rpm for 10 min. After washing twice with 250 μL of Exosome Immobilizing/Washing Buffer, the beads were added to the samples. To bind EVs against the beads, the mixture was agitated at 1,000 rpm for 2 h, followed by washing thrice with 250 μL of Exosome Immobilizing/Washing Buffer. Finally, bead-captured EVs were eluted with 100 mM Tris-HCl (pH 8.0) and 20 mM NaCl containing 4% SDS.

### Protein digestion

The protein lysate, purified EVs, and biotinylated BSA were treated with 20 mM tris(2-carboxyethyl) phosphine at 80°C for 10 min and alkylated using 35 mM iodoacetamide at room temperature for 30 min while being protected from light and subjected to clean up and digestion with SP3 using Maelstrom 8 Autostage. Briefly, two types of SeraMag SpeedBead carboxylate-modified magnetic particles (hydrophilic particles: CAT# 45152105050250 and hydrophobic particles: CAT# 65152105050250; Cytiva, Marlborough, MA, USA) were used. The beads were combined in a 1:1 (v/v) ratio, washed twice with distilled water, and reconstituted in distilled water at a concentration of 8 μg solids/μL. Consequently, 20 μL of reconstituted beads was added to the alkylated protein sample followed by 99.5% ethyl alcohol to bring the final concentration to 75% (v/v), with mixing for 5 min. The supernatant was discarded, and the pellet was washed twice with 80% ethyl alcohol. The beads were then resuspended in 80 μL of 50 mM Tris-HCl (pH 8.0) or 50 mM Tris-HCl (pH 8.0) containing 0.02% LMNG, followed by 500 ng of trypsin/Lys-C Mix (CAT# V5072, Promega, Madison, WI, USA) and mixed gently at 37°C overnight to digest proteins. The digested sample was acidified with 20 μL of 5% trifluoroacetic acid (TFA) and then sonicated with Bioruptor II (CosmoBio) at a high level for 5 min at room temperature. Samples were desalted using an SDB-STAGE tip (CAT# 7820-11200, GL Sciences Inc., Tokyo, Japan) or Evotip Pure (CAT# EV2015; EVOSEP, Odense, Denmark). The SDB-STAGE tip was washed with 25 μL of 80% ACN in 0.1% TFA, followed by equilibration using 50 μL of 3% ACN in 0.1% TFA. Then, the sample was loaded onto tip, washed using 80 μL of 3% ACN in 0.1% TFA, and eluted with 50 μL of 50% ACN in 0.1% TFA or 36% ACN in 0.1% TFA. The eluate was then dried using a centrifugal evaporator (miVac Duo Concentrator). The dried sample was redissolving in 8 μL of 2% ACN containing 0.1% TFA or 0.01% DMNG and transferred to the normal vial. Four microliters of the sample were injected into the LC-MS/MS system. Evotip Pure was used according to the manufacturer’s protocol. In brief, tip was washed with 20 μL of ACN in 0.1% formic acid (FA), followed by equilibration using 20 μL of 0.1% FA in H_2_O. Then, the sample was loaded onto tip and washed using 20 μL of 0.1% FA in H_2_O. Tip was filled with 0.1% FA in H_2_O until LC−MS/MS measurements.

### CoIP and on-beads digestion

IP lysis buffer [Pierce IP Lysis Buffer (CAT# 87788, Thermo Fisher Scientific, Waltham, MA, USA) containing protease inhibitors (CAT# 5892791001, cOmplete ULTRA Tablets, Sigma-Aldrich, MO, USA) and phosphatase inhibitors (CAT# 4906837001, PhosSTOP Tablets, Sigma-Aldrich)] was added to HeLa cells and mixed at 4°C for 30 min. The cell lysate was then centrifuged at 18,000 × *g* and 4°C for 30 min, and the supernatant was collected. Protein concentration in the protein extract was determined using a BCA protein assay kit (CAT# 23225, Thermo Fisher Scientific) and adjusted to 2 μg/μL with the IP lysis buffer. Co-immunoprecipitation (IP) of the HeLa cell lysate was performed using an automated Maelstrom 8 Autostage. Beads for coIP used Sera-Mag SpeedBeads Protein A/G Magnetic Particles (CAT# 17152104010150, Cytiva, Uppsala, Sweden), and antibodies used anti-RELA (NF-κB p65) antibody (CAT# ab16502, Abcam, Cambridge, MA, USA). To prepare the beads for immunoprecipitation, 1.5 μL of bead slurry was washed once with 500 μL of tris buffered saline containing 0.05% Tween20 (TBST). Then, the anti-RELA antibody was captured on the beads by mixing the beads at room temperature for 30 min in 200 μL of TBST containing 4 μg anti-RELA antibody and washing twice in 500 μL of TBST to remove the unbound antibody. Subsequently, 200 μL of HeLa cell lysate (2 μg/uL) was added to the beads and incubated 60 min at room temperature with end-over mixing, followed by washing thrice with 500 μL of the IP lysis buffer and twice with 500 μL of 50 mM Tris-HCl (pH 8.0). The beads were treated with 100 μL of 50 mM Tris-HCl (pH 8.0) or 50 mM Tris-HCl (pH 8.0) containing 0.02% LMNG, followed by 500 ng of trypsin/Lys-C Mix (CAT# V5072, Promega), and mixed gently at 37°C overnight to digest proteins. The beads were then aggregated from the digested sample using a magnetic stand (EpiMag HT (96-Well) Magnetic Separator; EpiGentek, Brooklyn, NY, USA) to collect the supernatant. The collected sample was treated with 20 mM tris (2-carboxyethyl) phosphine at 80°C for 10 min and alkylated using 35 mM iodoacetamide at room temperature for 30 min while being protected from light. Subsequently, the alkylated sample was acidified with 10 μL of 10% TFA (total volume 120 μL) and desalted using SDB-STAGE tip (elution: 36% ACN in 0.1% TFA), followed by drying in a centrifugal evaporator (miVac Duo Concentrator). The dried sample was redissolved in 10 μL of 2% ACN containing 0.1% TFA or 0.01% DMNG and transferred to normal vials (Thermo Fisher Scientific). One microliter of each sample was injected into the LC-MS/MS system.

### Sample preparation for SCP

As a collection plate for FACS-sorted cells, 10 µL of 2 ng/µL trypsin/Lys-C Mix (CAT# V5072, Promega) in 0.3% acetic acid was added to a 96-well plate (CAT# 0030128664, Eppendorf, Hamburg, Germany), sealed with plate seal (CAT# 5010-21951, GL Sciences Inc., Tokyo, Japan), and stocked at -80°C.

FreeStyle HEK293-F cells were cultured in DMEM medium supplemented with 10% FCS until they reached 80% confluence. Cells were collected, pelleted by centrifugation, and washed thrice with phosphate buffered saline (PBS). For FACS cell sorting, 5 µL DAPI was added to the 5 mL single-cell solution, and cell sorting was performed on the DAPI-negative live cell population with BD FACSMelody cell sorter. Single or ten cells were sorted into the collection plates, brought to room temperature, sealed with plate seal (CAT# 5010-21951, GL Sciences Inc.), and frozen at 80°C until use. Single or ten cells in 96-well plates were brought to room temperature and centrifuged at 500 × *g* for 3 min. Five microliters of 500 mM Tris-HCl pH 8.0 with 0.08% LMNG or 500 mM Tris-HCl pH8.0 was added to the plate, subjected to water bath-type sonication (Bioruptor II, CosmoBio, Tokyo, Japan) for 5 min, and mixed gently at 37°C overnight to digest proteins. The plates were centrifuged at 500 × *g* for 3 mins and 5 µL of 5% TFA was added and then treated with Evotip Pure according to the manufacturer’s protocol.

### DDA-MS by Typical LC-MS/MS

The redissolved peptides were directly injected onto a 75 μm × 12 cm nanoLC nano-capillary column (Nikkyo Technos Co., Ltd., Tokyo, Japan) at 50°C and then separated with a 30-min gradient (A = 0.1% FA in water, B = 0.1% FA in 80% ACN) consisting of 0 min 5% B, 30 min 45% B at a flow rate of 300 nL/min using an UltiMate 3000 RSLCnano LC system. The peptides eluted from the column were analyzed on an Orbitrap Exploris 480 (Thermo Fisher Scientific) using the InSpIon system (AMR) (Kawashima *et al*, 2023). MS1 spectra were collected in the range of 380–1,240 m/z at a 60,000 resolution to set an AGC targets of 3 × 10^6^ and a maximum injection time of 100 ms. The 50 most intense ions with charge states of 2+ to 5+ that exceeded 8.0 × 10^3^ were fragmented by collision induced dissociation with a normalized collision energy of 28%. MS2 spectra were collected in the range of more than 200 m/z at a 15,000 resolution to set an AGC targets of 1 × 10^5^ and a maximum injection time of “Auto.” The dynamic exclusion time was set to 30 s.

### DIA-MS by Typical nanoLC-MS/MS

The redissolved peptides were directly injected onto a 75 μm × 12 cm nanoLC column (Nikkyo Technos Co., Ltd., Tokyo, Japan) at 50°C and then separated with a 60-min gradient (A = 0.1% FA in water, B = 0.1% FA in 80% ACN) consisting of 0 min 8% B, 50 min 35% B, 57 min 70% B, 60 min 70% B at a flow rate of 200 nL/min using an UltiMate 3000 RSLCnano LC system. The peptides eluted from the column were analyzed using an Orbitrap Exploris 480 with an InSpIon system. MS1 spectra were collected in the range of 495–905 m/z at a 15,000 resolution to set an AGC targets of 3 × 10^6^ and a maximum injection time of 23 ms. MS2 spectra were collected at more than 200 m/z at a 30,000 resolution to set an AGC targets of 3 × 10^6^, a maximum injection time of “Auto,” and a normalized collision energy of 26%. The isolation width for MS2 was set to 8 m/z, and for the 500–900 m/z window pattern, an optimized window arrangement was used in Xcalibur 4.4 (Thermo Fisher Scientific).

### EVOSEP ONE LC-MS/MS

The Evotip Pure sample was analyzed using an EVOSEP ONE system (EVOSEP) equipped with an Orbitrap Exploris 480 mass spectrometer and InSpIon system. EVOSEP ONE was acquired using the Whisper 80 SPD method (gradient running time, 15 min; flow rate, 100 nL/min). The digested peptides were separated using an Aurora Rapid 75 C18 capillary column (5 cm × 75 μm i.d., particle size 1.7 μm; IonOpticks, VIC, Australia) at 60°C. Mobile phases A and B consisted of 0.1% FA in H_2_O and 0.1% FA in ACN, respectively. The peptides eluted from the column were analyzed using an Orbitrap Exploris 480 instrument with DIA. DIA parameters were slightly modified from a previously reported high-throughput proteome analysis (Ishikawa *et al*, 2022). MS1 spectra were collected in the range of 645–775 m/z at a 7,500 resolution to set an AGC targets of 1 × 10^6^ and a maximum injection time of 10 ms. MS2 spectra were collected at 200– 1800 m/z at a 30,000 resolution to set an AGC targets of 3 × 10^6^, a maximum injection time of “Auto,” and a normalized collision energy of 28%. The isolation width for MS2 was set to 8 m/z, and for the 650–770 m/z window pattern, an optimized window arrangement was used in Xcalibur 4.4 (Thermo Fisher Scientific).

To optimize DIA using the Whisper 80SPD method for smaller samples, we investigated the proteome coverage of the 49 DIA methods using 1 ng of cell digest. These DIA methods were designed using different combinations of precursor mass ranges (m/z 120, 180, and 240), mass resolutions (30,000, 45,000, and 60,000), and isolation window widths (8, 12, 15, 20, and 24 Da). In all methods, MS1 parameters were set as follows: mass resolution, 7,500; AGC targets, 1 × 10^6^; and maximum injection time, 10 ms. In DIA, the maximum injection time values at mass resolutions of 30,000, 45,000, and 60,000 were set at 55, 87, and 119 ms, respectively. AGC targets for MS2 was set to 3 × 10^6^ at 28% normalized collision energy.

### Protein identification and quantitative analysis from MS data

DDA-MS files were searched against the human protein sequence UniProt database (proteome ID UP000005640, 20,591 entries, downloaded on May 5, 2023) using Proteome Discoverer 3.0, with Sequest HT and Percolator (Thermo Fisher Scientific). The setting parameters were as follows: enzyme, trypsin; maximum missed cleavage sites, 2; precursor mass tolerance, 10 ppm; fragment mass tolerance, 0.02 Da; static modification, cysteine carbamidomethylation; and dynamic modification, methionine oxidation. DIA-MS files were also searched against an *in silico* human spectral library using DIA-NN (version: 1.8.1, https://github.com/vdemichev/DiaNN) (Demichev *et al*, 2020). First, a spectral library was generated from the human protein sequence UniProt database using DIA-NN. The Parameters for generating the spectral library were as follows: digestion enzyme, trypsin; missed cleavages, 1; peptide length range, 7–45; precursor charge range, 2–4; precursor m/z range, 395–1005; and fragment ion m/z range, 200–1800. “FASTA digest for library-free search/library generation;” “deep learning-based spectra, RTs, and IMs prediction;” “n-term M excision;” and “C carbamidomethylation” were enabled. The DIA-NN search parameters were as follows: mass accuracy, 10 ppm (15 ppm with Evosep One); MS1 accuracy, 10 ppm (15 ppm with Evosep One); protein inference, genes; neural network classifies, single-pass mode; quantification strategy, robust LC (high precision); and Cross-run normalization, Off. “Unrelated runs,” “use isotopologues,” “heuristic protein inference,” and “no shared spectra” were enabled. The MBR was turned off when searching for the identification of proteins and precursors and turned on for quantitative analysis. The protein identification threshold was set at 1% or less for both precursor and protein FDRs.

### Phosphoproteomics

100 µg of peptide digest was dissolved in 80% ACN in 0.1% TFA. Dissolved peptides were added to Fe-NTA Magnetic Agarose (80 µL of suspension used; Cat # A52284, Thermo Fisher Scientific) washed with 80% ACN in 0.1% TFA and mixed for 60 min at room temperature, followed by washing twice with 1 mL of the 80% ACN in 0.1% TFA and once with 1 mL of 0.1% TFA. The beads were treated with 200 μL of 3% polyphosphoric acid in 0.1% TFA or 3% polyphosphoric acid in 0.1% TFA containing 0.02% LMNG and then mixed for 10 min at room temperature to elute the phosphopeptides. The eluted phosphopeptides were desalted by SDB-STAGE tip (elution: 36% ACN in 0.1% TFA). The dried sample was redissolving in 10 μL of 0.01% DMNG and transferred to a normal vial. The redissolved peptides were directly injected onto a 75 μm × 30 cm nanoLC column (CAT# HEB07503001718IWF, CoAnn Technologies, Richland, WA, USA) at 60°C and then separated with a 100-min gradient (A = 0.1% FA in water, B = 0.1% FA in 80% ACN) consisting of 0 min 5% B, 86 min 35% B, 93 min 70% B, 100 min and 70% B at a flow rate of 150 nL/min using an UltiMate 3000 RSLCnano LC system. The peptides eluted from the column were analyzed on an Orbitrap Exploris 480 using the InSpIon system. MS1 spectra were collected in the range of 400–1,500 m/z at a 60,000 resolution to set an AGC targets of 3 × 10^6^ and a maximum injection time of “Auto.” The 50 most intense ions with charge states of 2+ to 4+ that exceeded 2.0 × 10^4^ were fragmented by collision induced dissociation with stepped normalized collision energies of 22%, 26%, and 30%. MS2 spectra were collected in the range of 200–1,800 m/z at a 30,000 resolution to set an AGC targets of 5 × 10^5^ and a maximum injection time of “Auto.” The dynamic exclusion time was set to 30 s. The MS files were searched against the human protein sequence UniProt database (proteome ID UP000005640, 20,591 entries, downloaded on March 7, 2023) using Proteome Discoverer 3.0 with Sequest HT and Percolator (Thermo Fisher Scientific). The setting parameters were as follows: enzyme, trypsin; maximum missed cleavage sites, 2; precursor mass tolerance, 10 ppm; fragment mass tolerance, 0.02 Da; static modification, cysteine carbamidomethylation; and dynamic modification, serine, threonine, and tyrosine phosphorylation. The protein identification threshold was set at 1% or less for both peptide and protein FDRs.

### LC-MS/MS analysis of biotinylated BSA

A mixture of 10 ng of biotinylated BSA digest and 5 µg of K562 cell digest (Promega) was added to streptavidin beads (10 µL of suspension used; Cat # 21152104010150, Cytiva), washed once with TBST, and mixed for 60 min at room temperature, followed by washing thrice with 1 mL of 50 mM Tris-HCl (pH 8.0) and 500 mM NaCl containing 0.5% SDS and once with 1 mL of TBS. The beads were treated with 50 μL of 8M guanidine-HCl (pH 1.5) in 0.5 mM biotin or 8 M guanidine-HCl (pH 1.5) in 0.5 mM biotin containing 0.02% LMNG and then mixed for 2 h at room temperature to elute the biotinylated peptides. The eluted peptides were desalted by SDB-STAGE tip (elution: 36% ACN in 0.1% TFA). The dried sample was redissolved in 10 μL of 0.01% DMNG and transferred to a normal vial. The redissolved peptides were directly injected onto a 75 μm × 12 cm nanoLC nano-capillary column (Nikkyo Technos Co., Ltd) at 50°C and then separated with a 30-min gradient (A = 0.1% FA in water, B = 0.1% FA in 80% ACN) consisting of 0 min 8% B, 26 min 45% B, 29 min 85% B, 30 min 85% B at a flow rate of 300 nL/min using an UltiMate 3000 RSLCnano LC system. The peptides eluted from the column were analyzed on Q Exactive HF-X using the InSpIon system. MS1 spectra were collected in the range of 450–1,500 m/z at a 120,000 resolution to set an AGC targets of 3 × 10^6^ and a maximum injection time of 100 ms. The 20 most intense ions with charge states of 2+ to 7+ that exceeded 8.0 × 10^3^ were fragmented by collision induced dissociation with normalized stepped collision energies of 22%, 26%, or 30%. The MS2 spectra were collected in the range of more than 200 m/z at a 60,000 resolution to set an AGC target of 2 × 10^5^ and a maximum injection time of 120 ms. The dynamic exclusion time was set to 30 s. The MS files were searched against the human protein sequence UniProt database (proteome ID UP000005640, 20,591 entries, downloaded on May 5, 2023) and BSA (UniProt Ac P02769) using PEAKS Studio 11 (Bioinformatics Solution Inc., Waterloo, Canada). The parameters were as follows: enzyme, trypsin; maximum number of missed cleavage sites, 4; precursor mass tolerance, 10 ppm; fragment mass tolerance, 0.02 Da; static modification, cysteine carbamidomethylation; and dynamic modification, lysine biotinylation. The protein identification threshold was set at 1% or less for both peptide and protein FDRs.

## Data analysis

Perseus v1.6.15.0 was used for principal component analysis (PCA) and Pearson’s correlation coefficient heatmap analysis with hierarchical clustering.

## Acknowledgments

This study was supported in part by JSPS KAKENHI under Grant Numbers 23H02465, 22KK0077, 20K20469, and 21K07877 and by AMED under Grant Number JP22ek0109586. We thank Hiroko Kinoshita and Yukiko Eifuku for their help with the reagents and sample preparation.

## Author Contributions

YK and OO conceived and designed the study. DN performed pretreatment for shotgun proteomics. YE performed FACS for SCP. MI conducted the LC–MS/MS measurements. RK performed the computational work. YK and RK wrote the manuscript, and MI, DN, and OO edited it. All the authors have read and approved the final version of the manuscript.

